# The interaction of relative age with maturation and body size in female handball talent selection based on only sport-specific criteria

**DOI:** 10.1101/2024.01.29.577867

**Authors:** Zsófia Tróznai, Katinka Utczás, Júlia Pápai, Gergely Pálinkás, Tamás Szabó, Leonidas Petridis

## Abstract

Talent identification is often affected by the relative age effects (RAEs) and/or maturation. Previous research has suggested that talent selection should include sport-specific technical tasks instead of body size and/or physical test measurements, assuming that the technical tasks are less influenced by interindividual variability in maturation. Our purpose was to examine the prevalence of RAEs and how biological maturity, body size, and body composition in relation to relative age affect talent selection among female handball players. Handball-specific drills and in-game performance were the criteria of a single talent selection program. Birth distribution of U14 female (*N* = 3198) handball players were analyzed. Body size, body composition, and bone age were assessed on 264 selected and 266 not-selected players. Body composition was assessed with InBody 720, whereas biological age from bone age. Chi-square was used to examine quartile distributions, and logistic regression to determine the effects of the predictors on the selection. In terms of all registered players, there was no difference in birth distribution. RAEs appeared at the first selection level and were evident at all levels. Quartiles differed only between the first and the last quartiles in body size and muscle mass. Only bone age differed between consecutive quartile or semi-year groups. Body size, body composition, and maturity had a significant, but of moderate power, effect on the selection. Larger body height increased selection odds by about 12%, larger muscle mass by 12% to 25%, larger percent body fat decreased selection odds by 7%, while larger bone age by 3.5-4 times. RAEs affected talent selection when applying sport-specific drills as selection criteria. Relative age was connected to bone age, but not convincingly to body size and muscle mass. Bone age had the largest effect on the selection, but this was not associated with larger body size or muscle mass.

## Introduction

To increase competitiveness in international competitions, talent identification programs are commonly adapted in youth sport aiming to select the most talented players within a single age group [1–3]. Selected players usually receive additional training and sport science support and have more competition opportunities [4] compared to non-selected players. It has been well documented, however, that talent selection at young ages is biased by the relative age effects (RAEs) and biological maturation [5–8]. As a result, relatively younger and/or less mature players are often excluded from selection, which may negatively affect their long-term athletic career [9–10]. The RAEs refer to birth inequalities between players born early and late within a single selection period (e.g., calendar year) [11]. Players born at the beginning of the selection period (relatively older) tend to be overrepresented compared to those born at the end of the selection period [12] (relatively younger).

Common explanations for the RAEs have focused on differences in maturation [13], body size [14], physical qualities [15–17], or a combination of all these [18]. Selection bias, particularly in team sports, may be further magnified when selection criteria are based primarily on measurements of body dimensions and/or physical tasks. Indeed, research in handball have revealed significant prevalence of the RAEs within selection systems suggesting unequal distribution in favor of relative older players [19–20].

To reduce the magnitude of RAEs and maturity, Cobley et al. [6] recommended that selection should include mainly technical and movement coordination skills. For instance, Vandendriessche et al. [21] highlighted that coordination tasks in soccer are not related to biological maturity. In a similar study with handball players, Matthys et al. [22] reported that sport-specific tasks are less influenced by body size and maturation than physical tests, thus their inclusion in talent selection could potentially separate talent identification from the variation in relative age and maturity status. Nevertheless, in a single talent selection including only sport-specific tasks, significant relative age effects were found among 13-14 years old female and 14-15 years old male handball players [23], which seem to challenge the potential of sport-specific tasks in reducing the RAEs.

Despite the available literature in talent development research, most findings rely on data from male athletes. A gender data gap seems to exist in this field with proportionally less data on female athletes compared to male athletes [24]. Given the continuously growing popularity of female sports, it is important to understand how the interaction of relative age with maturation and body size may affect talent development and the pathway across elite sports in a cohort of only female athletes separately from male athletes. Such data could improve the quality and efficiency of talent development systems designed more specifically for female athletes.

Within a single talent selection program which included only sport-specific tasks, the purpose of this study was to examine how maturity status, body size, and body composition in relation to relative age affect selection chances among female handball players. Based on the generally accepted assumption, we expected that relatively older players would be biologically more matured and would exhibit larger body dimensions and muscle mass compared to the relatively younger players, which would benefit them in the selection.

## Material and methods

### Experimental design

This study included a single, multi-level talent selection program for female handball players. To examine the RAEs prevalence before the selection we compared the birth distribution of all registered handball players with that of the corresponding average population. To quantify the RAEs during talent selection we examined the players’ birth distribution at every selection stage. Then, at each selection stage, body size, body composition, and biological maturity in relation to relative age and between selected and not-selected players were analyzed. Finally, using regression analysis we examined the effects of body size, body composition, and biological age (predictors) on the selection (dependent variable). Measurements were completed from November 1, 2021 until March 31, 2022. The study was approved by the Institutional Research Ethics Committee (Approval number: TE-KEB/23/2021).

### The selection program

The selection program consisted of three selection stages, starting from the local (club) level to the national level. An overall illustration of the selection program is presented in Fig 1.

**Fig 1.**
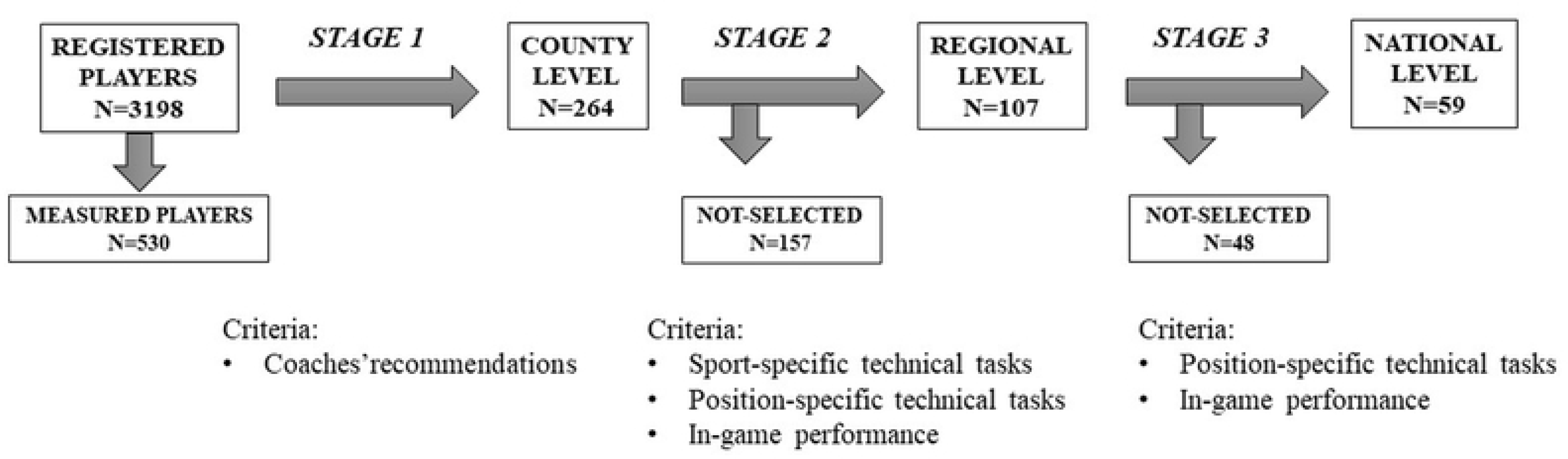
Flowchart of the Selection Program.

The selection stages were:

Local (club) level: coaches and staff members of all handball clubs and teams recommended players to participate in the Selection Program.

County level: selected players from the local level participated at this selection level, which included handball-specific generic skills, position-specific technical drills, and in-game performance. Detailed description of the selection tasks has been reported elsewhere [23]. Briefly, the handball-specific skills consisted of two tasks, the same for all players, one for defensive and one for offensive skills. For the defensive task, the players performed defensive footwork as quickly as they can between 9 cones placed in a zigzag line. For the offensive task, players had to perform a dribbling-shooting task two times consecutively, first dribbling the ball in a straight line and then in a zigzag line ending both times with a jump shot on goal. Points were awarded for the time results for the footwork task (maximum 5 points) and for the time duration (4 points) and the success of the shots (1 point) for the dribbling–shooting task. Both tasks were repeated two times with a short rest in-between. Position-specific drills included technical tasks according to playing positions: two tasks for backcourts (maximum 10–10 points), two for the wings (maximum 10–10 points), one for the pivots (maximum 20 points), and one for goalkeepers (maximum 20 points). Selection coaches evaluated passing and shooting accuracy, goal scoring, and technical execution. In-game performance evaluation included technique and efficacy of offensive and defensive movements (5–5 points, respectively). The players with the highest scores were selected for the next (regional) level.

Regional level: Players performed only the position-specific tasks, but under time pressure. At this level there was no rest between attempts; the tasks had to be performed at a higher intensity, thereby, increasing the level of difficulty. Evaluation and scoring were the same as at county level. In-game performance was again evaluated using the same criteria and scoring as at county level. Players with the highest scores were selected for the next (national) level.

National level: At this level, the evaluation included only in-game performance aiming to select the players for the age group national team. The selection for the national team was not examined in this study.

### Participants

Birth data of all registered female handball players born in 2008–2009 (*N*=3198) and birth data of the corresponding Hungarian population (*N*=95203) were obtained after permission from the Hungarian Handball Federation and from the Hungarian Central Statistical Office, respectively. Measurements were performed in a sample of *n*=530 players (birth date from 1 January 2008 to 31 December 2009). The players had to be registered members of handball clubs and had to hold a valid license to participate in official regional or national competitions. The measured players represented ∼17% of the total handball players in this bi-annual age group covering all competition levels and regions. The sample consisted of *n*=264 players (mean age: 13.1±0.6 years), who were selected to participate in the National Selection Program and of *n*=266 players (mean age: 13.2 ±0.6 years), who were not selected. Before the tests, the players and their parents received written information about the type and risk of the measurements and then the parents/guardians gave written consent for their children to participate. Consent was required by the ethics committee to provide the ethical approval for the study. We divided the players into eight groups based on their date of birth in quarter-year intervals (from Q1 to Q8). Frequencies of the quartile groups are presented in Table 1.

**Table 1.** Frequencies by relative age group across all measured levels.

|  |  |  | BASIC LEVEL |  | STAGE 1 |  | STAGE 2 |  | STAGE 3 |  |
| --- | --- | --- | --- | --- | --- | --- | --- | --- | --- | --- |
| Birth year |  | Average population | Registered players | Measured players | Not-selected | Selected | Not-selected | Selected | Not-selected | Selected |
| 2008 | Q1 | 11695 | 406 | 79 | 27 | 52 | 23 | 29 | 11 | 18 |
| 2008 | Q2 | 11664 | 386 | 68 | 30 | 38 | 19 | 19 | 5 | 14 |
| 2008 | Q3 | 13000 | 424 | 80 | 27 | 53 | 25 | 28 | 12 | 16 |
| 2008 | Q4 | 11967 | 351 | 53 | 24 | 29 | 21 | 8 | 4 | 4 |
| 2009 | Q5 | 11508 | 424 | 62 | 37 | 25 | 21 | 4 | 1 | 3 |
| 2009 | Q6 | 11428 | 409 | 79 | 52 | 27 | 17 | 10 | 6 | 4 |
| 2009 | Q7 | 12578 | 422 | 60 | 37 | 23 | 18 | 5 | 5 | 0 |
| 2009 | Q8 | 11363 | 376 | 49 | 32 | 17 | 13 | 4 | 4 | 0 |
| **Total** |  | **95203** | **3198** | **530** | **266** | **264** | **157** | **107** | **48** | **59** |

Q1-Q8: quartiles groups two years period. (2008: Q1: Jan.-Mar.; Q2: Apr.-Jun.; Q3: Jul.-Sep.; Q4: Oct.-Dec.) (2009: Q5: Jan.-Mar.; Q6: Apr.-Jun.; Q7: Jul.-Sep.; Q8: Oct.-Dec.)

### Anthropometry, body composition, and biological status

Prior to performing the selection tasks, the players took part in anthropometric, body composition, and bone age measurements. Due to injury or technical limitations, the bone age of 21 players and the body composition of 2 players was not measured. Anthropometric measurements were taken based on the recommendations of the International Biological Program (IBP) [25]. Body height was measured with an anthropometer (DKSH Switzerland Ltd, Zurich, Switzerland) to the nearest millimeter. Test-rest reliability for body height was ICC= 0.99, 95%CI (0.998-1.000). Body mass and body composition were determined using Inbody 720 (Biospace Co., Seoul, Korea) bioimpedance device. The athletes were measured in the morning hours wearing shorts and t-shirts. Participants were instructed not to consume food or drink two hours and not to perform strenuous physical activity 24 hours before the measurements. Percent body fat and muscle mass were processed. Test-retest reliability for body mass was: ICC= 1.00, 95%CI (1.000-1.000); for skeletal muscle mass: ICC= 0.99, 95%CI (0.998-1.000); and for percent body fat: ICC= 0.99, 95%CI (0.996-1.000).

Biological maturity was estimated based on bone age using an ultrasound-based device (Sunlight BoneAge, Sunlight Medical Ltd, Tel Aviv, Israel). This method estimates the stage of skeletal development and has been found reliable in boys up to 16 and in girls up to 15 years [26]. Measurements were performed at the wrist region of the left hand. Participants placed their arm on a horizontal surface between the transducers. The transducers were aligned with the growth zone of the forearm (radius and ulna) at the junction of the distal epiphysis and diaphysis. Then, in the initial position, the transducer was attached to the forearm at a pressure of approximately 500 g and emitted 750 kHz ultrasound at the measurement site for each measurement cycle. One measurement cycle lasted approximately 20 seconds and was repeated five times. The instrument estimated bone age (in years and months) using equations based on the speed of ultrasound (SOS) and the distance between the transducers. The difference between chronological age (CA) and bone age (BA) was used to estimate the maturity status of the participants. Test-retest reliability of bone age estimation was: ICC= 0.98, 95%CI (0.942-0.992).

### Statistical analysis

Frequencies were used to present distribution by relative age group. Data from anthropometric, body compositions, and bone age measurements were checked for outliers and for normal distribution (Kolmogorov-Smirnov test). Data of 29 athletes were excluded (Tukey method). Test-retest reliability was checked using absolute agreement 2-way mixed-effects model intraclass correlation coefficient (ICC). Differences in observed and expected distributions were tested using chi-square (*χ^2^*) test. Intergroup differences between Q1 to Q8 groups were tested with one-way analysis of variance (partial *η*^2^ effect size) using Tukey post hoc test with Bonferroni correction for multiple comparisons or with the Kruskal-Wallis test when normality was violated. Differences between the selected and not-selected groups were examined using independent sample *t*-test (Cohen *d* effect size) or Mann-Whitney *U*-test when normality was violated. Finally, binary logistic regression was used to determine the effects of body size, body composition, and biological age (predictors) on the selection (dependent variable). The odds of selection (0=not-selected, 1=selected) were examined in separate regression models according to body size (body height and body mass), body composition (skeletal muscle mass and percent body fat), and biological status (bone age and maturity status). IBM SPSS 25.0 was used in statistical analysis; significance was set at *p*<.05.

## Results

Differences in birth distribution between the registered handball players and the average population were not significant (*χ^2^*=12.6; *p* =.081). Significant differences in relative age distribution were observed at all levels of selection when compared to the distribution of the registered players (county level: *χ^2^*=31.3; *p*<.001, regional level: *χ^2^*=50.6; *p*<.001, and national level: *χ^2^*=44.1; *p*<.001). Between consecutive selection levels RAEs showed a marginally non-significant increase from county to regional level (*χ^2^*=13.9; *p* =.052) and a non-significant increase from regional to national level (*χ^2^*=6.8; *p*=.451). At the last (national) level there was no player born in the last two quartiles (Q7 and Q8) (Fig 2). Kolmogorov-Smirnov tests showed that the distributions of body height, skeletal muscle mass, and maturity status were normally distributed, whereas body mass, percent body fat, and bone age deviated from normal distribution.

**Fig 2.**
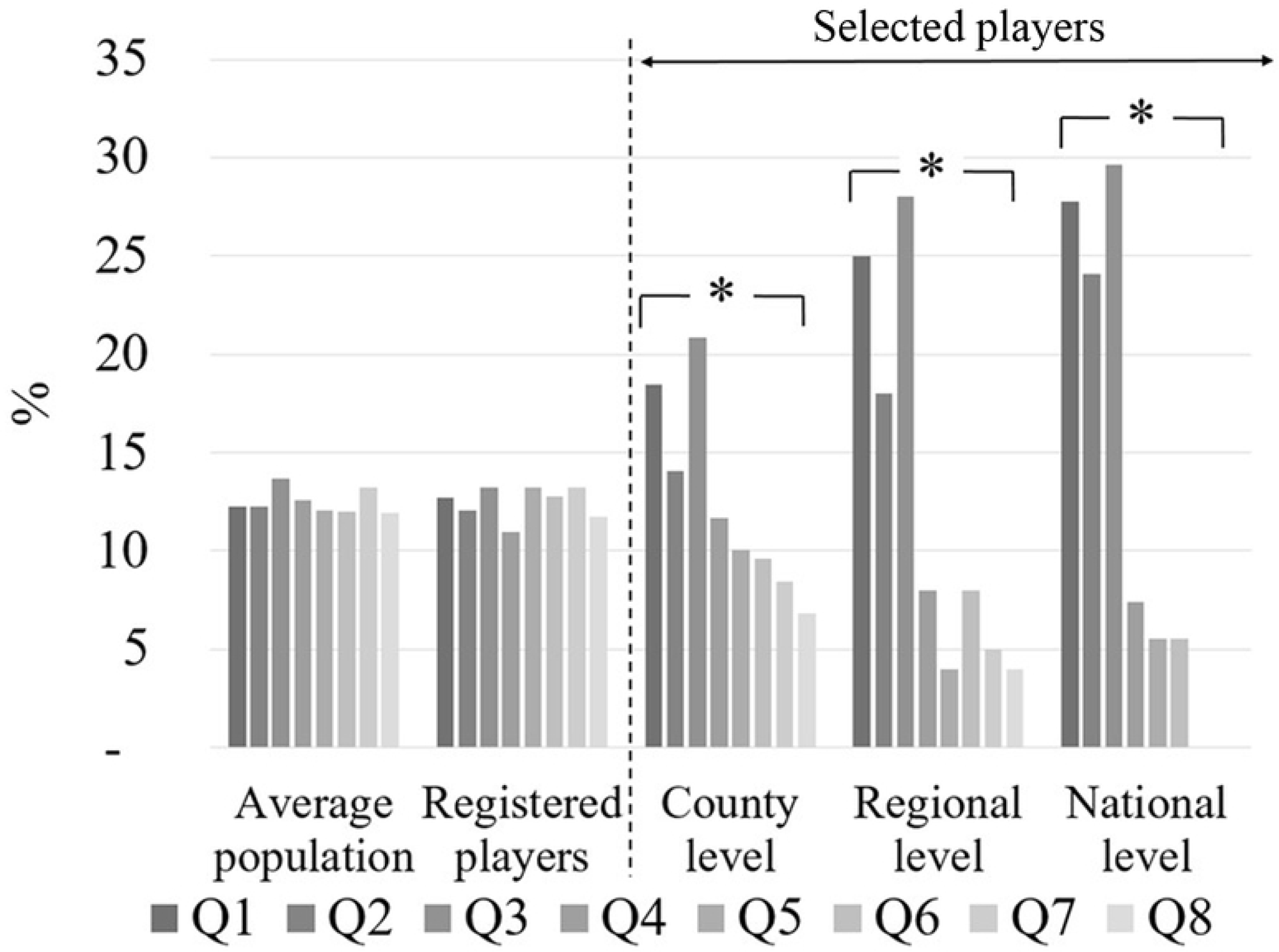
Birth distribution of the average population, of all registered players in the examined age group, and of the selection program. Q1-Q8: quartiles groups two years period (2008: Q1: Jan.-Mar.; Q2: Apr.-Jun.; Q3: Jul.-Sep.; Q4: Oct.-Dec.) (2009: Q5: Jan.-Mar.; Q6: Apr.-Jun.; Q7: Jul.-Sep.; Q8: Oct.-Dec.)*: *p*<.05 compared to registered players.

Boxplots of the measured variables in relation to the players’ relative age are presented in Fig 3. Differences were found only between the first (Q1, Q2) and the last quartiles (Q7, Q8) for body mass (Fig 3A) (H_501(7)_=20.55 *p* =.005), body height (Fig 3B) (F_(7)501_=3.68, *p* <.001, *η_p_*^2^=0.05), skeletal muscle mass (Fig 3D) (F_(7)499_=4.90, *p* <.001, *η_p_* ^2^=0.065), bone age (Fig 3E) (H =144.36 *p* <.001) and biological maturity (Fig 3F) (F_(7)481_=3.40, *p* =.001, *η_p_*^2^=0.048). Percent body fat did not differ between relative age groups (Fig 3C). Only bone age differed between consecutive quartile or semi-year groups (Fig 3E).

**Fig 3.**
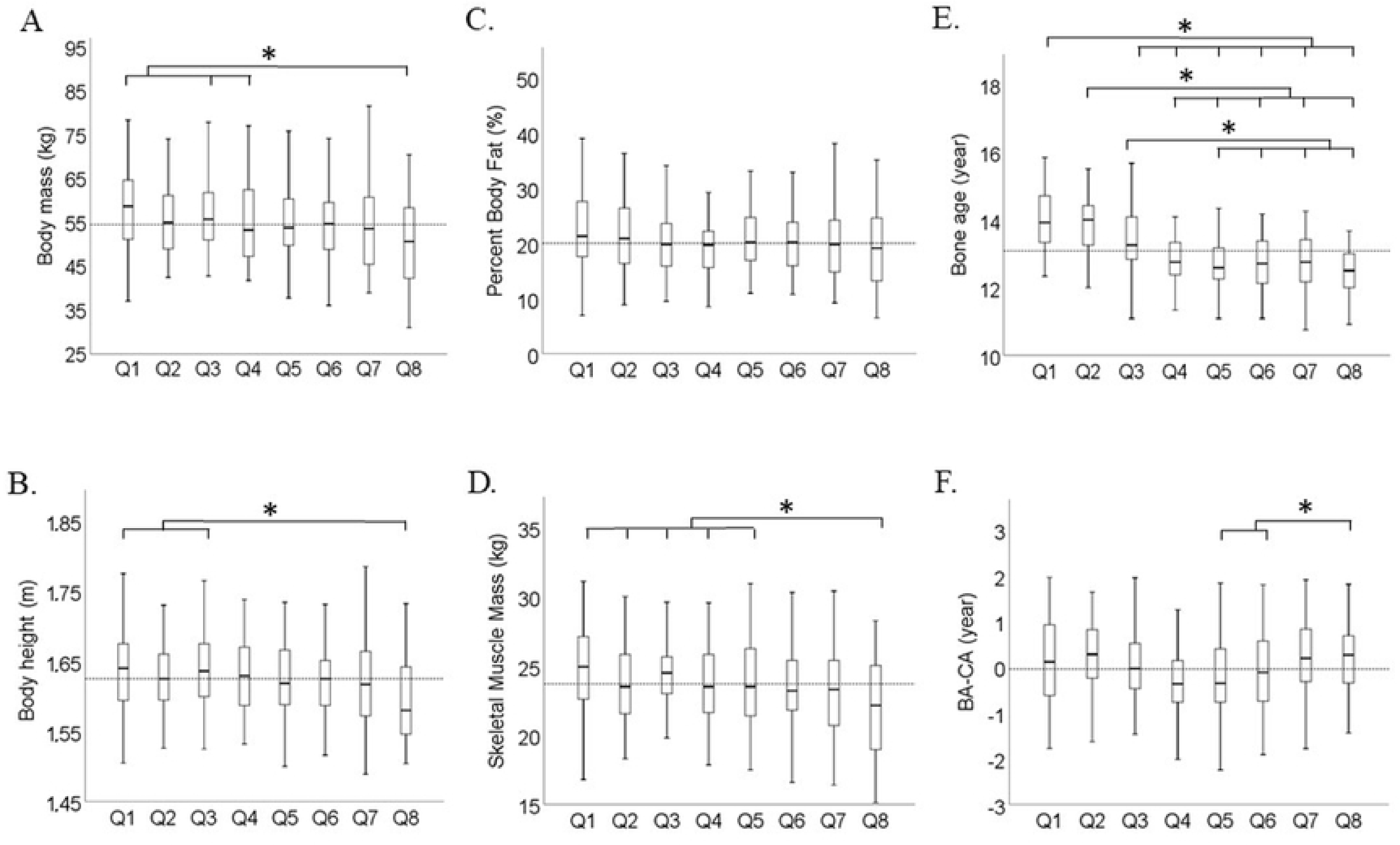
Boxplots of all measured players by relative age group for (A) body mass, (B) body height, (C) percent body fat, (D) muscle mass, (E) bone age and (F) maturity status (Bone age-Chronological age). The dashed line represents overall mean value. Q1-Q8: quartiles groups two years period. (2008: Q1: Jan.-Mar.; Q2: Apr.-Jun.; Q3: Jul.-Sep.; Q4: Oct.-Dec.) (2009: Q5: Jan.-Mar.; Q6: Apr.-Jun.; Q7: Jul.-Sep.; Q8: Oct.-Dec.)*: *p*<.05 between quartiles.

Fig 4 summarizes the results for the examined variables between selected and not-selected players at each selection level. Differences were evident at the first and second selection stages, however effect size of at least medium magnitude (Cohen *d*>0.5) were found only for maturation status at county stage (*d*=0.51, 95% CI: 0.33-0.64), and for body height and skeletal muscle mass at the regional stage (*d*=0.58, 95% CI: 0.33-0.84 and *d*=0.56, 95% CI: 0.30-0.82, respectively). Selected and not-selected players did not differ in any of the examined variable at the national selection stage.

**Fig 4.**
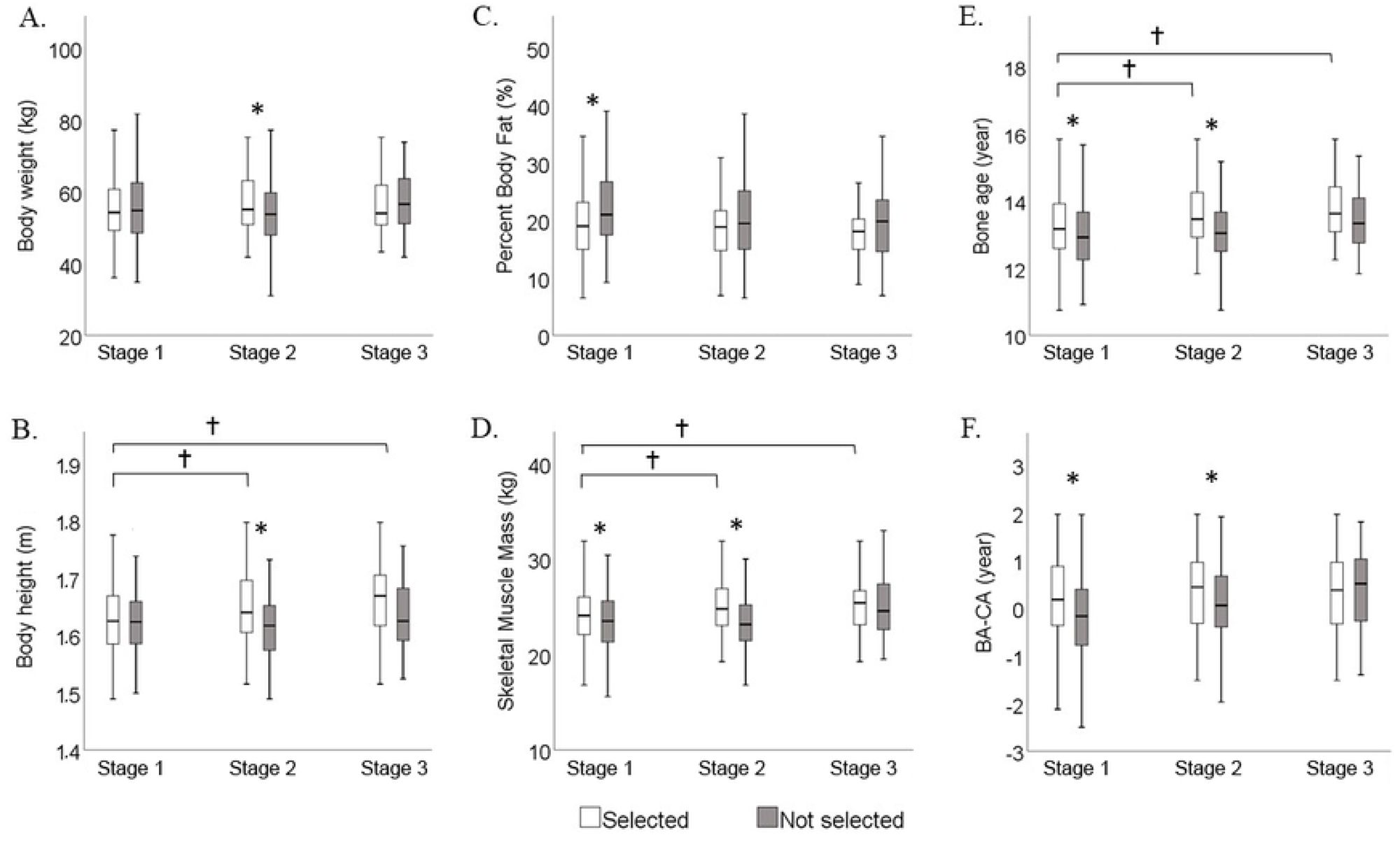
Boxplots of the selected and not-selected players at each selection stages for (A) body mass, (B) body height, (C) percent body fat, (D) muscle mass, (E) bone age and (F) maturity status (Bone age-Chronological age). *: *p*<.05 between selected and not-selected players, ⩾:p<.05 between selected players at different selection stages.

Tables 2-4 show the results of the binary logistic regression models for body size, body composition, and biological maturity, respectively. At basic level we did not include bone age measurements in the regression analysis, because not-selected players were measured 2-4 months later than selected players potentially resulting in biased analysis. At this selection level, body size had no effect on the selection. Body composition components contributed significantly with an explanatory power of about 8% (Table 2). Larger skeletal muscle mass increased selection odds by about 12% for every one kilogram, while larger percent body fat decreased selection odds by 7% for every one percent of body fat.

**Table 2.** The results of the binary logistic regression at basic level (Stage 1) (dependent variable: selected/not-selected).

| Models | Variables | B | Wald | *p* | Expl(B) | 95%CI |
| --- | --- | --- | --- | --- | --- | --- |
| $\chi^2$ 2.95 | Body mass | -0.02 | 1.95 | .162 | 0.98 | 0.96-1.01 |
| *p* .229 | Body height | 0.03 | 2.59 | .107 | 1.03 | 0.99-1.07 |
| **R<sup>2</sup>** 0.01 | Constant | -3.99 | 2.15 | .142 | 0.02 |  |
| $\chi^2$ 29.43 | Skeletal muscle mass | 0.12 | 12.86 | <.001 | 1.12 | 1.05-1.19 |
| *p* <.001 | Percent body fat | -0.07 | 22.31 | <.001 | 0.93 | 0.90-0.96 |
| **R<sup>2</sup>** 0.08 | Constant | -1.25 | 2.94 | .086 | 0.29 |  |

At county level all three models were significant (Table 3). Biological maturity explained about 15%, body size about 10%, and body composition about 13% of the variance in selection. Bone age had the largest effect on selection odds; one year increase in bone age increased selection odds by 3.5 times. In addition, one centimeter increase in body height and one kilogram increase in muscle mass increased selection odds by about 12% and 25% respectively.

**Table 3.**
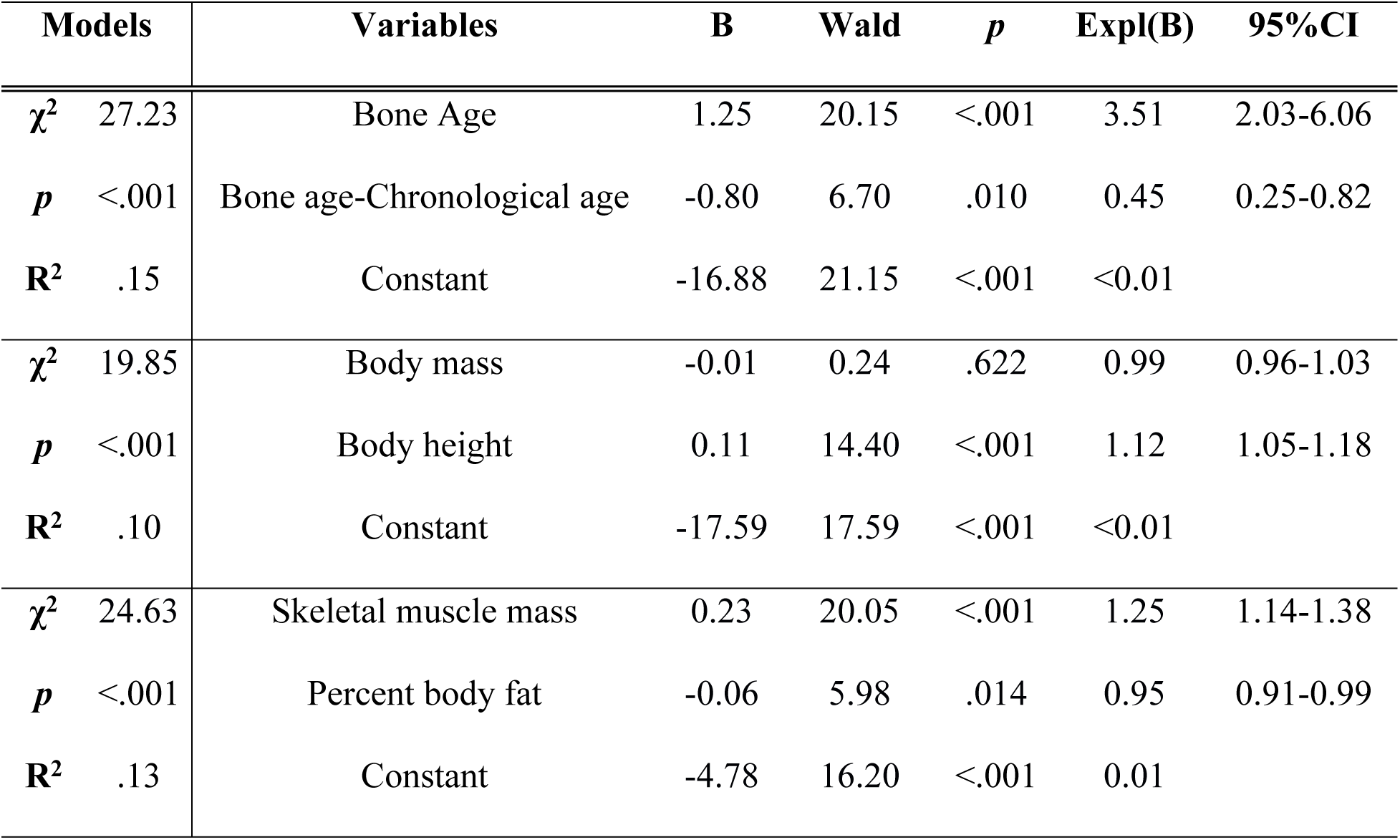
The results of the binary logistic regression at county level (Stage 2) (dependent variable: selected/not-selected).

| Models | Variables | B | Wald | *p* | Expl(B) | 95%CI |
| --- | --- | --- | --- | --- | --- | --- |
| $\chi^2$ 27.23 | Bone Age | 1.25 | 20.15 | <.001 | 3.51 | 2.03-6.06 |
| *p* <.001 | Bone age-Chronological age | -0.80 | 6.70 | .010 | 0.45 | 0.25-0.82 |
| **R<sup>2</sup>** .15 | Constant | -16.88 | 21.15 | <.001 | <0.01 |  |
| $\chi^2$ 19.85 | Body mass | -0.01 | 0.24 | .622 | 0.99 | 0.96-1.03 |
| *p* <.001 | Body height | 0.11 | 14.40 | <.001 | 1.12 | 1.05-1.18 |
| **R<sup>2</sup>** .10 | Constant | -17.59 | 17.59 | <.001 | <0.01 |  |
| $\chi^2$ 24.63 | Skeletal muscle mass | 0.23 | 20.05 | <.001 | 1.25 | 1.14-1.38 |
| *p* <.001 | Percent body fat | -0.06 | 5.98 | .014 | 0.95 | 0.91-0.99 |
| **R<sup>2</sup>** .13 | Constant | -4.78 | 16.20 | <.001 | 0.01 |  |

At regional level the explanatory power of the three models decreased, significant effect was found for biological maturity and body size (Table 4). One year increase in bone age increased selection odds by more than 4 times, whereas one centimeter increase in body height by about 10%.

**Table 4.**
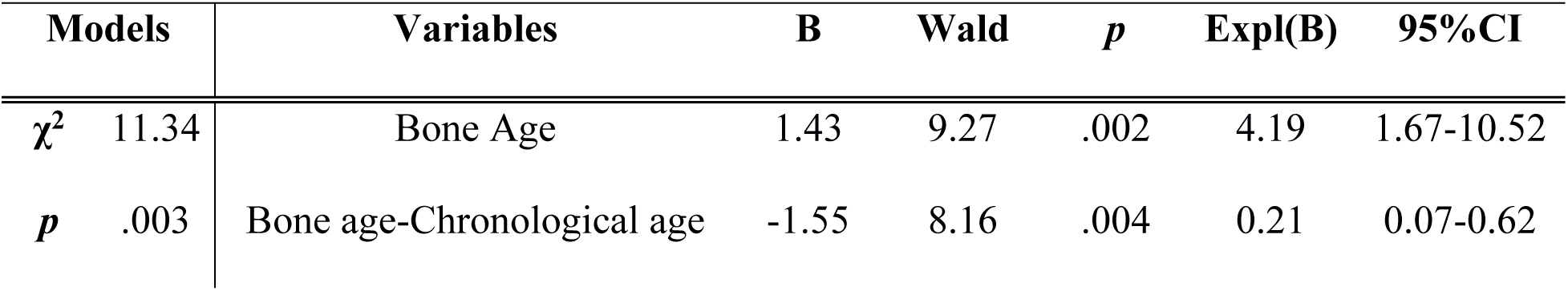
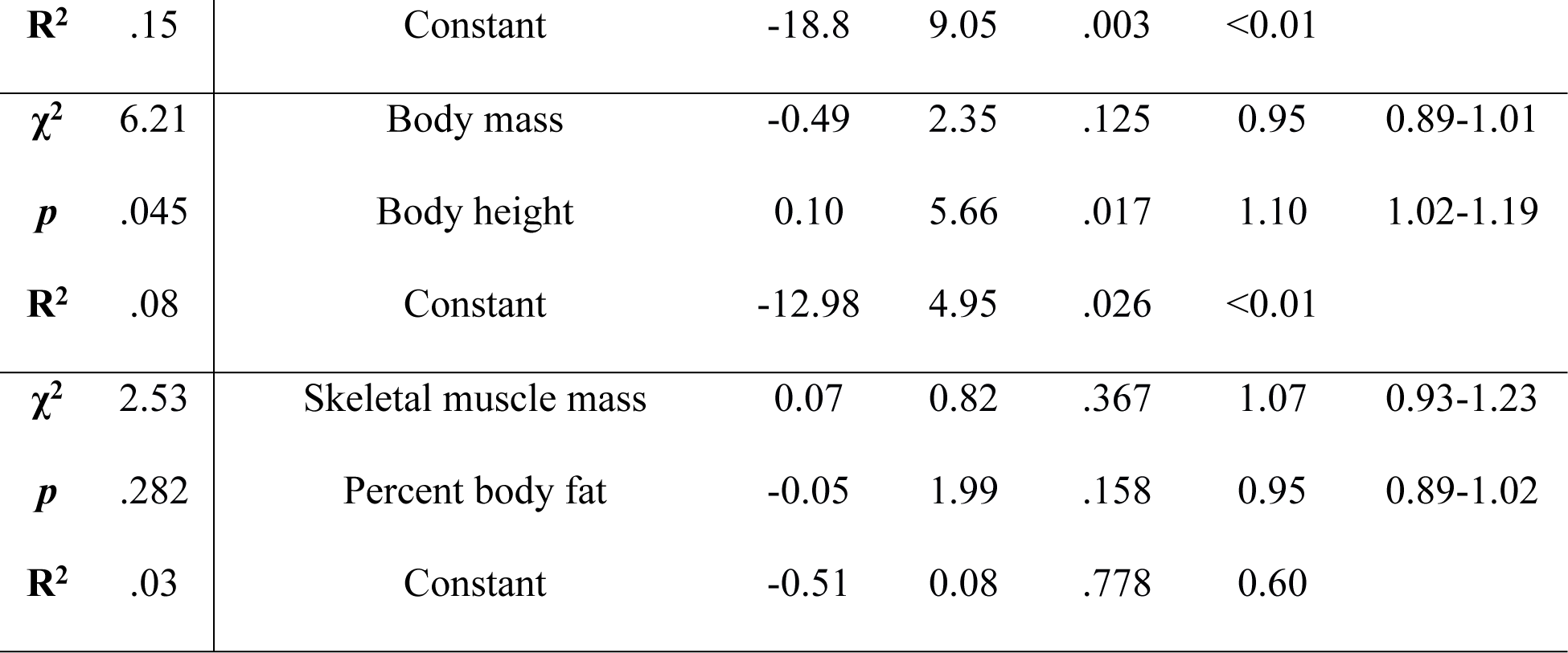
The results of the binary logistic regression at regional level (Stage 3) (dependent variable: selected/not-selected).

## Discussion

The purpose of this study was to investigate the interaction of relative age with body size, body composition, and biological maturity within a single selection program among adolescent female handball players. In general, selected players were taller, had larger muscle mass, less percent body fat, and were biologically more developed than not-selected players, which is not surprising considering the importance of physical qualities in handball [24–30]. However, the connection of these characteristics to the relative age was not straightforward.

Talent selection and the progression to the elite stages during an athletic career have been identified as facilitators in the prevalence of the RAEs [6,8,31]. To reduce the impact of the RAEs, previous research [6,32] has suggested to use only sport-specific tasks as selection criteria because the latter seem to be independent from biological development [22]. Yet, the results revealed significant RAEs at all selection levels. Considering that birth distribution at the initial level of all registered players in this age group did not differ from that of the average population, the results indicate that the RAEs initiated during the selection process. These results are in accordance with previous findings [23], which examined the prevalence of RAEs in talent selection program of similar structure, indicating a systemic effect of relative age even in this type of talent selection. It should be noted however, that the RAEs were evident only within the bi-annual age grouping, quartiles within one-year intervals did not differ significantly. Bi-annual age grouping is a common practice in international and domestic competitions; however, it seems that during talent selection the two-years age categories may increase the magnitude of the RAEs reinforcing the need to consider alternative grouping strategies.

Typically, relatively older players exhibit larger body size and more advanced maturity compared to the relatively younger players gaining in this way significant performance benefits during the selection. To test this hypothesis, we compared these measures according to relative age groups. The results did not convincingly confirm this assumption. Based on the entire sample (including both selected and not-selected players), differences between relative age groups were sporadic and inconsistent. In addition, significant differences were evident only between the first and the last quartile groups (e.g., in body height and muscle mass), that is, between players of above 1.5 years difference in chronological age. Bone age, as an indicator of biological development, was the only measure, which showed more consistent differences across relative age with relatively older players having higher values compared to their relative younger peers. Importantly, however, more advanced bone age was not definitely associated with larger body dimensions and muscle mass.

Another relevant factor is the maturity status, which is expressed as the difference between biological and chronological age. The distribution of maturing status was similar between quartile groups suggesting that prior to the selection, the proportion of early/on-time/late maturing players follows a normal pattern in all relative age groups. In other words, relative younger groups did not significantly exhibit higher proportion of early maturing players, which could benefit them in the selection. Together with the bone age data, the results indicate that the differences in biological development are simply attributed to quarter- or semi-year differences in chronological age, but not to variations in maturity status.

Interestingly, despite the differences in bone age between relatively younger and older players body dimensions did not differ. A possible reason is that even before the selection process, the body height and body mass of the players was around the 75^th^ percentile of the normative data for the average population (1.63 m and 55 kg respectively) [33] suggesting that handball attracts girls who are genetically predisposed to large body dimensions independently of their relative age and maturation. Noteworthy, at the time of the measurements most of the examined girls were after their menarche and after their age at peak height velocity, likely indicating reduced influence of biological maturity on body size development.

A second aim was to examine the effects of anthropometric factors and maturation on the selection for each selection stage separately. Since differences between relative age groups were limited and to make the analysis more concise, we grouped the players only based on the outcome of the selection, but not according to their relative age.

At the first stage, body composition components and biological age along with maturity status appeared as significant predictors in the selection. This is an interesting finding considering that the selection at this level was based solely on the subjective evaluation of team coaches. This confirms earlier reports suggesting that physicality and advanced maturity influence the coaches’ eye even when these attributes are not selection criteria. For example, Krahenbühl and Leonardo [34] reported that among U14 male and female handball players, coaches perceived relatively older players as better players than relatively younger ones with the former having more competition time. The logistic regression results revealed a significant, but of moderate explanatory power contribution of muscle mass and percent body fat in the selection. Larger skeletal muscle mass increased selection odds by about 12%. Interestingly, body size had no effect on the selection at this stage, as this was demonstrated by the differences between selected and not-selected players and by the logistic regression results.

At the second stage, selection included sport-specific and playing position-specific tasks. Although these tasks required mainly technical skills, the contribution of body height, muscle mass, and biological age was significant with their explanatory power being the largest at this level (Table 3). This is confirmed also by the significant differences between selected and not-selected players (Fig 4) favoring those with larger body height and muscle mass and more advanced maturity. Sport-specific technical tasks have been suggested to be affected by biological maturation to a limited extent, thus potentially may reduce selection bias [22], although there was a study where maturity had a positive, but small, contribution to the variation in soccer-specific technical skills [35]. Our findings collectively suggest that the technique-based selection tasks cannot eliminate the impact of relative age and consequently of biological development on the selection among female handball players. The large selection odds of bone age in the logistic regression model (almost four times increase with increase in bone age; Table 4) clearly highlight the impact of biological development on the selection, which most probably is not limited only to the well-known benefits in body size and physical qualities, but likely affects several other key elements in handball, like coordination, movement control, or game intelligence [36]. It has been suggested previously [13] that the selection chances of relatively younger players may increase with advanced maturity, however this was not clearly supported from our findings. Early maturation did not increase selection odds, even more it seemed to negatively affect the selection (Tables 3 and 4). A possible explanation for this discrepancy is that during puberty, advanced maturity in girls intensifies gains in body mass accompanied also by increases in body fat mass and percent body fat [37]. Such changes in body composition, however, may impair speed, agility, and eventually athletic performance [38].

The last selection corresponds to the elite stage of the selection. The selected players at this level represented less than 2% of all players in this age group. Differences between the selected and not-selected players were limited, which possibly indicates that players of different body size and body composition characteristics were excluded earlier during the selection forming in this way a more homogenous group. At this level only bone age had a significant effect on the selection increasing selection odds, which may provide further explanation of the occurrence of RAEs considering the strong connection of relative age and bone age. Compared to normative data from the average population, the mean body height of the elite players (1.66±0.07 m) was between the 75-90^th^ percentile [33] and it was larger by 5.6 cm than the mean value of the average population demonstrating the superior body size characteristics of the elite handball players even at a young age. On average, muscle mass of the elite players was larger by 1.5 kg and percent body fat lower by 2.6% compared to the entire sample. Body height and body mass of the elite female handball players is comparable with that of the Greek 13.8-year-old female preliminary national handball team [27] (1.66±0.07 m; 57.3±7.8 kg) and with the selected U14 players from Croatia (1.66±0.07 m; 57.0±7.4 kg) [39].

The analysis is limited by playing positions. Talent selection was completed according to the playing position of the players, which due to the different anthropometric profile and body composition characteristics may increase variability between players (particularly between backcourts and wings) and therefore may affect the statistical analysis. In addition, the effects of the included variables on the selection were examined only comprehensively, but not according to the selection tasks. Considering the different nature of each task, the analysis may yield effects of various magnitude and/or direction for each task separately.

## Conclusion

Overall, the findings have demonstrated the occurrence of the RAEs in a single talent selection program, which included only sport-specific technical tasks. Findings suggest (1) that relative age was connected to bone age, but not convincingly to body size and muscle mass (2) small effects of the examined variables on the selection, which may not provide strong explanation for the occurrence of the RAEs. However, the small differences between relatively older and younger players for each variable separately are added likely resulting in notable advantage for the former and (3) that further factors (not discussed here) may account also for the variability in the selection. This indicates, that the RAEs cannot be simply explained with differences in maturation, body size, or muscle mass, but rather constitute a more complex phenomenon.

## Acknowledgments

We thank all the athletes, who participated in the measurements as well as their coaches and their parents/guardians for giving their consent. We are grateful for the assistance received from the Hungarian Handball Federation, which allowed this study to be carried out.

